# Low immune response after 1.5 years of primary SARS-CoV-2 infection and Covishield vaccination lead to SARS-CoV-2 reinfection

**DOI:** 10.1101/2022.05.12.491584

**Authors:** Anita M. Shete, Deepak Y. Patil, Rima R. Sahay, Gajanan N. Sapkal, Gururaj R. Deshpande, Pragya D. Yadav

**Affiliations:** Indian Council of Medical Research-National Institute of Virology, Pune, India Pin-411021

## Abstract

We have investigated six COVID recovered cases with two doses of Covishield vaccination followed by reinfection. The primary SARS-CoV-2 infection found to occur with B.1 and reinfection with Omicron BA.1 and BA.2 variants. The genomic characterization and duration between two infections confirms these cases as SARS-CoV-2 reinfection. The mutation analysis of the reinfection cases correlated with immune evasion potential of BA.1 and BA.2 sub lineages. The immune response determined at different time intervals demonstrated boost post two dose vaccination, decline in pre-reinfection sera post 7 months and rise post reinfection. Apparently, these cases suffered from SARS-CoV-2 reinfection with the declined hybrid immunity acquired from primary infection and two dose covishield vaccination. This suggests the need for booster dose of vaccination. Besides this, multiple non-pharmaceutical interventions should be used to cope up with SARS-CoV-2 infection.

---

Here, we have investigated six individuals with primary SARS-CoV-2 infection with two doses of Covishield vaccination followed by reinfection.

The cases acquired primary SARS-CoV-2 infection during the first wave during March 2020 to October 2021 (first wave). The cases were then vaccinated with two doses of Covishield vaccine post three months of primary infection. They were followed until the occurrence of reinfection during January 2022 to February 2022 (third wave) in India. Suspected reinfection cases were identified as a positive SARS-CoV-2 Real time RT-PCR (Ct value >35) ≥90 days after the initial positive test and at least one consecutive negative test result between two incidences [1]. The naso/oropharyngeal swab and blood specimens were collected at four different time points namely primary SARS-Cov-2 infection, post 60 days of second vaccine dose, pre-reinfection (post 7 months of second dose), reinfection (post 10 months of second dose). The SARS-CoV-2 confirmation and genomic characterization was done using E gene specific Real time RT-PCR [2] and next generation sequencing [3]. The IgG immune response was determined using SARS-CoV-2 S1-RBD specific IgG ELISA [4]. We have also measured the neutralizing antibody (NAb) titres of sera against Delta, Omicron and ancestral SARS-CoV-2 B.1 variant using a plaque reduction neutralization test (PRNT) [5].

During primary infection, all the six cases tested positive for SARS-CoV-2 using Real time RT-PCR (rRT-PCR) specific for E gene (viral RNA copy number 8.9×10^3^ to 1.6×10^6^/ml). The next generation sequencing of the six cases revealed the infecting SARS-CoV-2 variant to be prototype B.1 strain (Fig. 1 A). Four cases were asymptomatic; while two symptomatic cases presented with moderate fever, sore-throat, dry cough, generalized weakness, loss of appetite, loss of smell and taste. The mean IgG antibody titre of the cases at reinfection was found to be 300 (Fig. 2 A). Besides this, the NAb geometric mean titre (GMT) of the sera against B.1, Delta and Omicron was 643 (95% CI 295-1404), 43 (95% CI 6-291), 1.9 (95% CI 0.16-22) respectively (Fig. 2 B-D).

**Figure 1:**
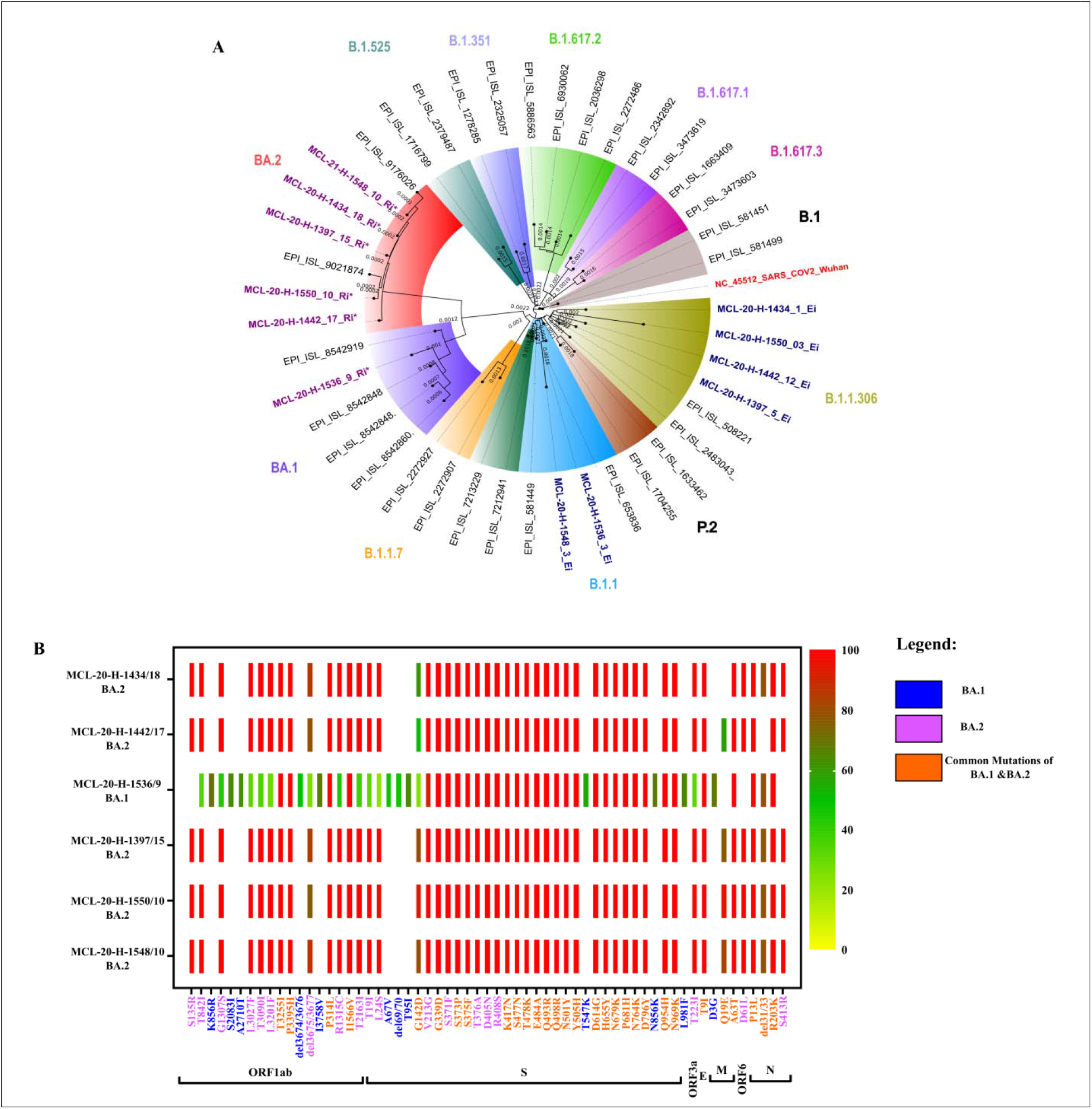
Phylogenetic and variant analysis of SARS-CoV-2 reinfection cases: A) Molecular phylogenetic tree designed based on a neighbor joining analysis of SARS-CoV-2 sequences retrieved from clinical specimens of cases during primary infection, reinfection and reference sequences with bootstrap⍰of 1000 replicates. The sequence of primary infection and reinfection cases are highlighted with blue color and violet color respectively. Reference sequences are marked with black color along with prototype Wuhan strain in red color. B) Heat map analysis of SARS-CoV-2 variants BA.1 and BA.2 and frequency differences between the mutations and deletions. Mutations in BA.1, BA.2 and common mutations in BA.1 & BA.2 are marked with blue, pink and orange color respectively.

**Figure 2:**
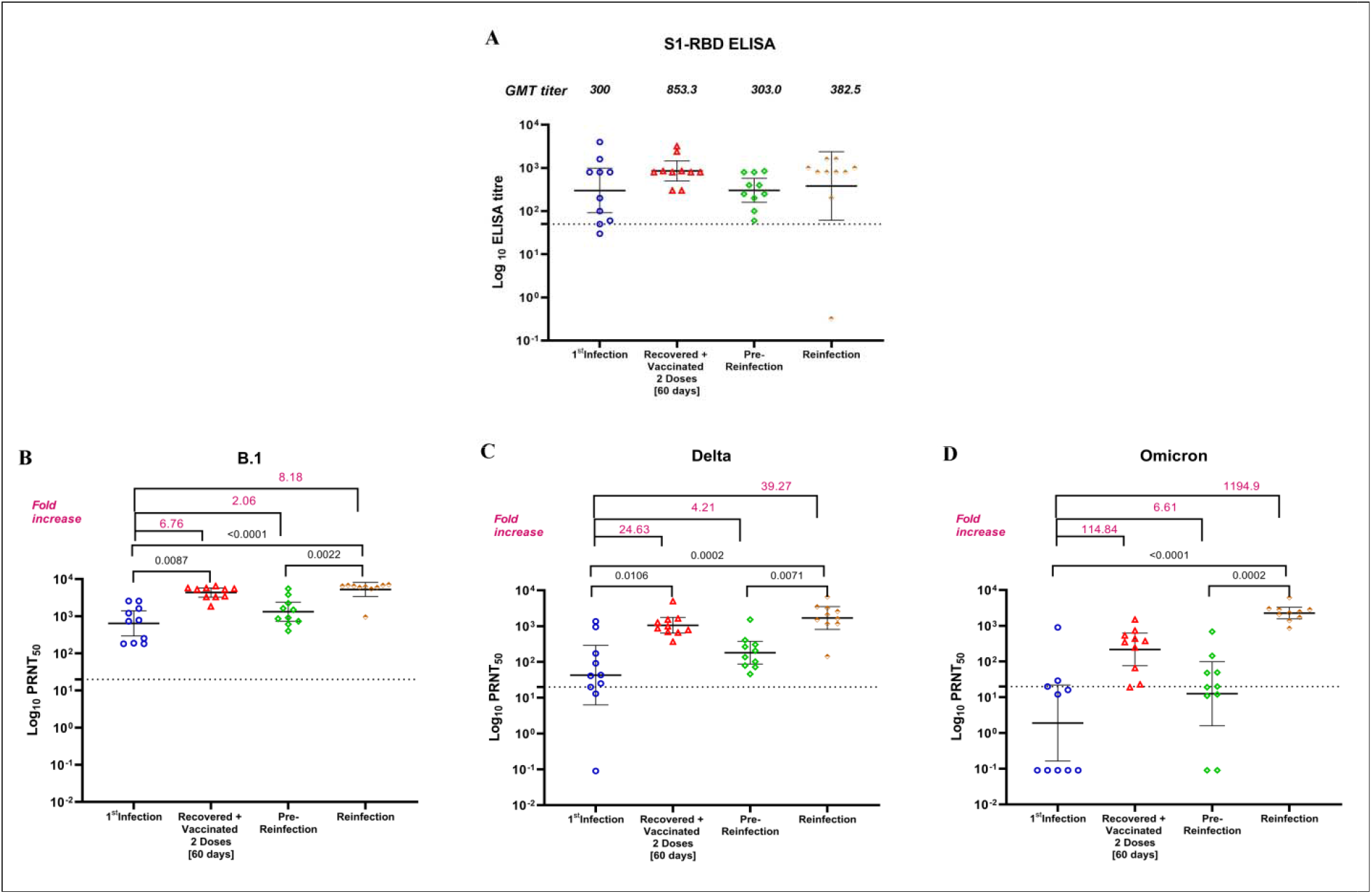
IgG and neutralizing antibody response at different time points during primary infection and reinfection: A) Anti-SARS-CoV-2 IgG antibody response determined with S1-RBD ELISA during primary infection, post sixty days of second dose vaccination, pre-reinfection and reinfection. Neutralizing antibody titres determined with plaque reduction neutralization test (PRNT) during primary infection, post sixty days of second dose vaccination, pre-reinfection and reinfection against B) B.1 variant C) Delta variant D) Omicron variant. The dotted line represents the limit of detection of the assay. Data are presented as mean values +/- standard deviation (SD).

The serum samples collected after 2 months from second dose vaccination shown 2.8-fold rise in IgG and 6.8, 24.5, 475-fold rise in NAb titres against B.1, Delta and Omicron respectively than the primary infection. The pre-reinfection sera collected after 7 months of second vaccination demonstrated reduction in IgG antibodies titre (2.8-fold) and also in the NAb titres against B.1 (3.3-fold), Delta (5.9-fold), and Omicron (17.3-fold) (Fig. 2 A-D).

The SARS-CoV-2 reinfection amongst these COVID-recovered two dose vaccinees found to occur after 10 months after second dose vaccination. Of these, three cases were asymptomatic; while three symptomatic cases had myalgia, hoarseness of voice, dry cough, cold and sore-throat. The genomic characterization of six clinical specimens (viral RNA copy number 1.1×10^5^ to 1.4×10^8^/ml) revealed infecting SARS-CoV-2 variant to be Omicron sub lineage BA.1 (n=1) and BA.2 (n=5) (Fig. 1 A). The genomic analysis and duration between two infections confirms these cases as SARS-CoV-2 reinfection. The mutation analysis of the reinfection cases represented the mutations and deletions in BA.1 (K856R, S2083I, A2710T, del3674/3676, I3758V, A67V, del69/70, T95I, T547K, N856K, L981F, D3G), BA.2 (S135R, T842I, G1307S, L3201F, del3675/3677, R1315C, T2163I, T19I, L24S, V213G, S371F, T376A, D405N, R408S, T223I, D61L, S413R) and common mutations/deletions of BA.1 and BA.2 (T3255I, P3395H, P314L, I1566V, G142D, G339D, S373P, S375F, K417N, S477N, T478K, E484A, Q493R, Q498R, N501Y, Y505H, D614G, H655Y, N679K, P681H, N764K, D796Y, Q954H, N969K, T9I, Q19E, A63T, P13L, del31/33, R203K) (Fig. 1 B). Of which, common mutations found in BA.1 & BA.2 (K417N, S477N, T478K, E484A, Q493R, Q498R, N501Y, Y505H) correlates with immune evasion potential of BA.1 and BA.2 sub lineages. The NAb GMT of reinfection cases sera against B.1, Delta and Omicron were found to be 5261 (95%CI: 3413-8111), 1685 (95%CI: 811-3501) and 2262 (95%CI: 1558-192.4) respectively (Fig. 2 B-D). These titres represent significant boost in immune response post reinfection compared to pre-reinfection sera. As the reinfection occurred with Omicron sub lineages, about 1000-fold rise was observed in NAb titres in reinfection sera with Omicron than B.1 and Delta. Apparently, IgG titre showed only 1.3-fold increase at reinfection (Fig. 2 A).

The findings of the study observed increasing immune response after two dose vaccination than at primary infection. Subsequently, this hybrid immunity found to decline to very low level at 7 month post second vaccination with which cases got reinfection. This suggests the need to protect the community through booster dose of vaccination and prevent further infections following personal hygiene and non-pharmaceutical interventions.

## Conflict of Interest Disclosures

No conflict of interest exists

## Funding

Financial support was provided by the Indian Council of Medical Research (ICMR), New Delhi at ICMR-National Institute of Virology, Pune under intramural funding ‘COVID-19’.

## Acknowledgement

Authors gratefully acknowledge the staff of ICMR-NIV, Pune including Dr. Rajlaxmi Jain, Mrs. Triparna Majumdar, Mrs. Savita Patil, Mr. Annasaheb Suryawanshi, Ms. Pranita Gawande, Ms. Jyoti Yemul, Mr. Yash Joshi, Ms. Manisha Dudhmal, Ms. Aasha Salunkhe and Mr. Chetan Patil, for extending excellent technical support.

## Notes

### Competing Interest Statement

The authors have declared no competing interest.

